# Prefrontal cortex dopamine responds to the total valence of stimuli

**DOI:** 10.1101/2024.12.11.627942

**Authors:** Yating Yang, Willem Parent, Hunter Rice, Risa Mark, Matthew Desimone, Monzilur Rahman, Ian T. Ellwood

## Abstract

The prefrontal cortex (PFC) dopamine system plays an essential role in cognitive flexibility, working memory and psychiatric disease, but determining the conditions under which dopamine in the PFC is released remains an open problem. Both rewarding and aversive stimuli have been found to trigger release, but studies have disagreed on whether the valence of a stimulus or other variables like novelty and salience are the most important. Here we report on recordings of dopamine-dependent fluorescence using a high-sensitivity dopamine indicator. We deliver an array of rewarding, aversive and mixed valence stimuli, as well as stimuli without any obvious valence. We observe that stimuli without valence, as well as the omission of expected stimuli, do not lead to large changes in fluorescence, even when these stimuli and omissions are both novel and engaging. In contrast, both rewarding and aversive stimuli lead to increases in fluorescence, with the most rewarding and most aversive stimuli leading to the largest increases. We test the effect of adding an aversive component to a rewarding stimulus and find that the increases in fluorescence are consistent with a summation of the rewarding and aversive components. We propose that dopamine release in the PFC responds to the total valence of a stimulus, in contrast with the traditional view of basal ganglia dopamine release that depends on the net valence.

## Introduction

The mesocortical dopamine system, which consists of dopaminergic fibers that project from the ventral tegmental area (VTA) and substantia nigra pars compacta (SNc) to the prefrontal cortex (PFC) (1), has several unique features, when compared with the mesolimbic system. Despite stimulation of the VTA being strongly reinforcing, selectively stimulating dopamine release in the PFC is neither reinforcing nor aversive in conditioned place preference tests and intracranial self-stimulation (2–4). Furthermore, aversive stimuli are particularly effective at eliciting PFC dopamine release, unlike in the NAc core and lateral shell, where release is inhibited. Recordings of PFC dopamine release using electrophysiology, calcium imaging, fast scan cyclic voltammetry and fluorescent dopamine indicators have all found that aversive stimuli trigger release (4–8), a property which may arise from the relatively larger number of excitatory inputs that PFC-projecting dopamine neurons receive from the lateral habenula (LHb) (9, 10). Nonetheless, PFC dopamine release is also triggered by rewards (3, 7, 8) and, when restricted to rewarding stimuli, can show reward prediction error-like qualities similar to the mesolimbic dopamine system (3), though this is not consistent across studies.

Given that PFC dopamine release occurs for both rewards and aversive stimuli, a natural question is what will happen when the valence of a stimulus is mixed, containing both rewarding and aversive components. In the mesolimbic dopamine system, this question has a simple answer. Dopamine release in the core and lateral shell regions of the nucleus accumbens, as well as the dorsomedial striatum, are well described by reward prediction error (RPE) theory (11, 12), especially in classical conditioning experiments. In RPE theory, dopamine release depends on stimuli only through their net valence, i.e. how good the stimuli are minus how bad they are. (See (13) for an exception to this rule.) In the language of RPE theory, net valence is typically called reward, and is denoted *R*, but this quantity can be negative for aversive stimuli and vanishes for stimuli that balance good and bad qualities. Net valence is a natural variable for reinforcement learning as it determines if an action is worth executing or an event is, on the balance, good or bad.

Since PFC dopamine is not reinforcing, and responds to both reward and aversion, net valence is not necessarily the right variable. Consider an experiment in which one delivers a sequence of stimuli, starting with a purely rewarding, unpredicted stimulus, and adds an increasingly large aversive component (Fig. 1A). During this process, the net valence of the stimulus will slowly decrease until it becomes zero and then becomes negative. In line with this, the amount of dopamine release in the NAc following the stimulus will decline and then become a dip in concentration once the stimulus is net aversive.

**Fig. 1.**
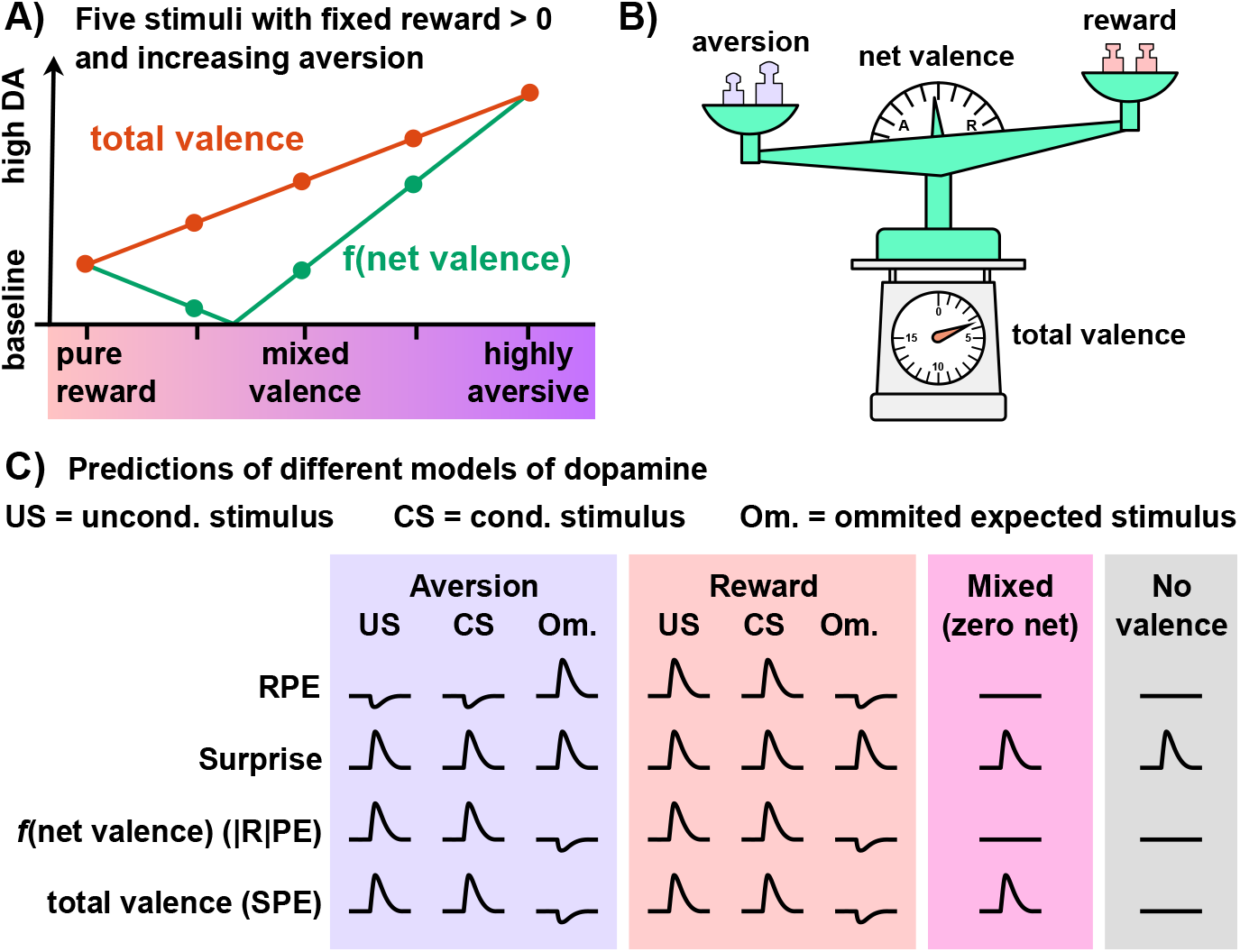
Comparing models of net valence, total valence and surprise. **A)** Given five stimuli, each of which has a non-zero rewarding component and a varying amount of aversion, we note that any model of net valence must predict zero dopamine release somewhere between a purely rewarding and highly aversive stimulus, which, in this toy example, is between the 2nd and 3rd stimuli (green line). A theory of total valence, on the other hand will interpolate between the two endpoints (orange line). **B)** Cartoon of the difference between a theory of net valence, which acts like a pan balance between reward and aversion, and total valence, which sums reward and aversion, here represented as a spring scale. **C)** Various theories of dopamine and their responses to unpredicted stimuli that are either aversive, rewarding, mixed valence or lacking any valence. The canonical RPE theory predicts inhibition by aversion and activation by reward, but treats a stimulus where reward and aversion are balanced as equivalent to no stimulus at all. If dopamine responds to surprise, it should respond to both reward, aversion and omissions of rewarding and aversive stimuli. As an example of a theory that depends on net valence, but predicts release for both reward and aversion, we illustrate |R|PE, which takes the absolute value of R before computing prediction errors. This theory predicts no release for a balanced mixed valence stimulus. Finally, a theory of total valence has similar predictions to |R|PE, but differs for balanced mixed valence stimuli, where it predicts release.

If we consider the effect of this same sequence of stimuli on the PFC dopamine system, it is unclear what should happen. Since both rewarding and aversive stimuli lead to release, we expect that the purely rewarding and highly aversive stimuli will induce dopamine release. What should happen between these two extremes is difficult to guess. One possibility is that, as the rewarding and aversive components of the event become balanced, the amount of dopamine release at the stimulus will become nearly zero (Fig. 1A, green). This would be consistent with PFC dopamine being a function of net valence. Even if the stimulus has strongly rewarding and aversive components, only their difference affects the PFC dopamine system so that a net neutral stimulus is equivalent to no stimulus at all.

In contrast, one might find that adding an aversive component to a rewarding stimulus will *not* lead to a reduction in dopamine release, but instead an *increase* and the amount of dopamine release will interpolate between what is seen with a pure reward and a reward mixed with a strongly aversive stimulus. This would show that PFC dopamine cannot be a function of net valence alone, but must depend on reward and aversion through more than just their difference. As we will suggest below, a natural model in this case is that PFC dopamine depends on the sum of rewarding and aversive components of a stimulus, which we will refer to as total valence (Fig. 1A, orange). If we think of net valance as a sort of pan balance that determines whether a stimulus leans toward aversion or reward, total valence is analogous to a spring scale placed under the pan balance, which measures the total weight placed on either side (Fig. 1B).

As a final observation, we note that when a system has an unsigned response to both valences, it is natural to question if valence is even the right variable to consider. Many dopamine neurons in the VTA respond to alerting sensory cues (14) or may respond more strongly to rare rewards than expected from RPE (15). Similarly, PFC dopamine neurons may respond to aspects of a stimulus like “novelty” or “salience”. Rather than being a signal that conveys how well or poorly recent events have gone, PFC dopamine might be an alerting signal that triggers PFC attention to recent events, both good and bad, whenever something unexpected happens. (See, for example, the recent proposal, (8)). This hypothesis would fit well with evidence that PFC dopamine increases attention to cues and enhances the signal to noise of PFC neural activity (2, 4).

While conceptually appealing, such hypotheses make strong predictions about dopamine release. If all prediction errors (i.e. surprises) lead to dopamine release, omitting an expected stimulus should induce dopamine release. Similarly, if PFC dopamine release is triggered by novelty, we should find that exploring a new environment should cause dopamine release, regardless of the valence of that environment. Finally, one should expect that there are stimuli with little or no apparent valence, but which are salient and engaging, that elicit large amounts of dopamine release.

An outline of the predictions of four models of dopamine is shown in Fig. 1C. We note several important features of these models, which should be compared with our experimental results below. First, RPE predicts that aversion will inhibit dopamine release and omissions of aversive stimuli will increase it. Second, a theory of dopamine based around surprise (i.e. all prediction errors) predicts release on both rewarding and aversive stimuli, but also on omissions of either. Third, any model that is only a function of net valence, such as | R | PE, which depends on net valence, R, through its absolute value, predicts that there will be no release when reward and aversion are balanced in a mixed valence stimulus. Finally, we consider total valence prediction error theory, *S*PE, which is described in more detail in the discussion. This model has identical predictions to | R | PE except for mixed valence stimuli, where it predicts that rewarding and aversive components add.

## Results

To record from PFC dopamine, we expressed a dopamine-dependent fluorophore GRAB_DA_3h, which has a high signal to noise and sensitivity (7, 16, 17). Fluorescence from the indicator was recorded with fiber photometry through an implanted optical fiber over the medial prefrontal cortex (mPFC) (18–20). Our implant locations, shown in Fig. S2A, were primarily in the prelimbic cortex (PL), though, due to the small size of PL, fluorescence from the infralimbic cortex likely contributed to our recordings as well. In some of our experiments, we contrast our mPFC recordings with recordings from the nucleus accumbens core (NAc core) region (Fig. S2A). For these recordings we used a lower sensitivity, but similarly high signal-to-noise indicator, GRAB_DA_3m (7).

### Conditioned aversive stimuli, but not omission of unconditioned stimuli elicit release in fear conditioning

The effects on PFC dopamine release of painful stimuli and cues associated with painful stimuli have been examined in previous studies (4, 6–8, 21). Here we show increases in fluorescence at a conditioned stimulus are due to a fear association by extinguishing the association and measuring the reduction in induced fluorescence. We also control for the effects of anxiety by using a non-conditioned cue and show that omitting an expected painful stimulus does not increase fluorescence.

Mice were trained in a classical, cued, delay fear conditioning paradigm (22, 23). The conditioning phase consisted of eight pairings of a conditioned auditory stimulus (CS) lasting five seconds that was terminated by an unconditioned painful stimulus (US), consisting of a 500 ms footshock. Semi-randomly interleaved with the CS, a 5 second, non-conditioned stimulus was also presented eight times, but was not terminated by the US. We note that we used a long intertrial interval of 2-3 minutes because preliminary recordings (data not shown) had revealed that the increase in fluorescence following a shock lasted over a minute. Mice showed both freezing and darting responses to the CS, which diminished with extinction (Fig. S2B,C)

We found, as expected from prior studies (4, 6–8), that the US elicited large increases in fluorescence (Fig. 2A,C). We observed that the CS also elicited an increase in fluorescence, even on the first trial when it had no association, but the evoked fluorescence rose with each pairing, more than doubling in peak magnitude by the last, eighth, pairing (Fig. 2C, I). The non-conditioned stimulus similarly elicited an increase in fluorescence on the first trial, and although the induced fluorescence slightly grew over the conditioning phase, the rise did not reach significance (pairwise *t*-test, *n* = 10, *t* = 1.89, *p* = 0.09) and was significantly smaller than what was observed for the CS (Fig. S2D, 2I).

**Fig. 2.**
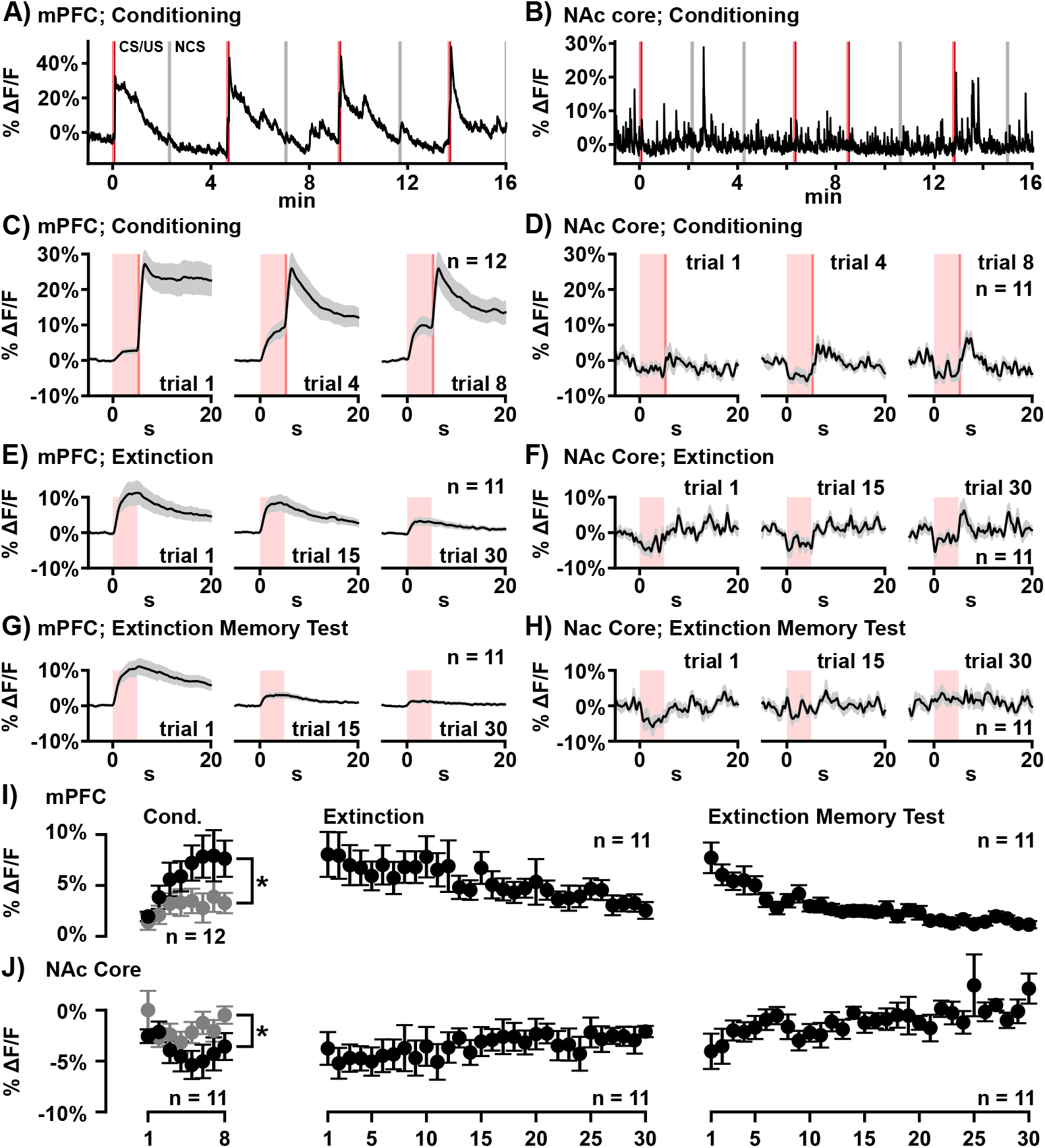
Conditioned aversive stimuli, but not omitted painful stimuli increase fluorescence. (* *p <* 0.05, ** *p <* 0.01). Error bars in this and subsequent figures represent standard error of the mean (s.e.m.) across animal subjects. For panels A-H, the left and right panels are recordings in the mPFC and NAc core respectively. **A-B)** Recording of fluorescence in a single animal in response to the first four CS/US pairs (red vertical lines) and the NCS (gray). **C-D)** Perievent average fluorescence during the CS and US during conditioning. Gray region represents s.e.m. **E-F)** Perievent average fluorescence during trials 1, 15, and 30 of extinction. **G-H)** Perievent average fluorescence during trials 1, 15, and 30 on a subsequent day test of extinction memory. **I)** Average across animals of ΔF*/*F during the five seconds of the CS. The increase in ΔF*/*F was significantly higher for the CS than the NCS (pairwise *t*-test, *t* = 2.69, *p* = 0.023). The average of the first five trials and last five trials were significantly different during extinction (pair-wise *t*-test, *t* = 2.87, *p* = 0.017) and the test of extinction memory (pair-wise *t*-test, *t* = 5.13, *p* = 0.0004 **J)** Same as panel I for recordings in the NAc core. The decrease in fluorescence for the CS was signficantly larger than for the NCS (pairwise *t*-test, *t* = −2.56, *p* = 0.029). The average of the first and last five trials were significantly different for both extinction (pairwise *t*-test, *t* = − 3.01, *p* = 0.013) and the test of extinction memory (pairwise *t*-test, *t* = −3.06, *p* = 0.012)

Following conditioning, the CS was presented in isolation 30 times. We did not observe any increase in fluorescence at the moment when the shock would have occurred on any of the extinction trials, including the first (Fig. 2E). Instead, fluorescence simply decayed back to baseline. We note that this finding was surprising to us, as we expected that a large positive reward prediction error (an aversive stimulus being unexpectedly omitted) would induce dopamine release.

Over the course of the thirty extinction trials, the CS associated dopamine release slowly decreased in tandem with the extinction of the fear association (Fig. 2I, S2B). We also recorded fluorescence on the next day in a test of extinction memory. We observed that fluorescence on the CS briefly returned to its peak post-conditioning magnitude, but subsequently quickly diminished, similar to the fear responses of the mice (Fig. 2G, I, and S2B).

We also performed recordings in the NAc core in this experiment. We found a near mirror image of the mPFC recordings with a CS-associated dip in fluorescence following conditioning (Fig. 2D) and a slow decrease in the dip depth during extinction (Fig. 2F, J) and the test of extinction memory (Fig. 2H, J). Although it is not the main focus of this study, we note that there was some variability in the responses of the NAc core recordings, a subset of which showed phasic increases in fluorescence at the shock, and, on average we find no change in fluorescence at the US. After extinction, we observed, on average, a small increase in dopamine following the CS (Fig. 2F, rightmost plot), which was not present in the mPFC recordings, though we did not observe a phasic response on the first omission trial as has been found in only some regions of the NAc (13, 24–27).

Examining Fig. 1C, blue region, the increases in mPFC fluorescence on both the US and CS, but not on the omission of the US, make it unlikely that either reward prediction error theory or a model that responds to all prediction errors can describe mPFC dopamine release. However, the recordings are still consistent with a model that depends on net valence, as long as its dependence is positive for aversive stimuli, and a theory of total valence. Note that, at least for the conditioned stimulus, the recordings in the NAc core are consistent with a reward prediction error theory.

Finally, we repeated our analysis for the isosbestic fluorescence, but did not find an increase in the fluorescence after conditioning nor a decrease in extinction memory, though we did find a small but significant rise in fluorescence during extinction for the mPFC mice (Fig. S2F); however, this change was much smaller than the decrease seen in the excitation fluorescence and in the opposite direction.

### Aversive but not safe novel environments elicit release in the elevated plus maze

To test the effects of innate fear and exploration on mPFC dopamine release, mice were run in a elevated plus maze (EPM) assay (28). We modified the standard protocol of the EPM by initially confining mice to right closed arm for 5 minutes by a partition. This allowed dopamine transients triggered by placing mice into the chamber to return to baseline and gave us an opportunity to compare fluorescence during exploration of the open arms with the unexplored left closed arm to control for any effects on mPFC dopamine release from exploration, as opposed to aversion. A single animal example, and an average over all recordings are shown in Fig. 3A, B.

**Fig. 3.**
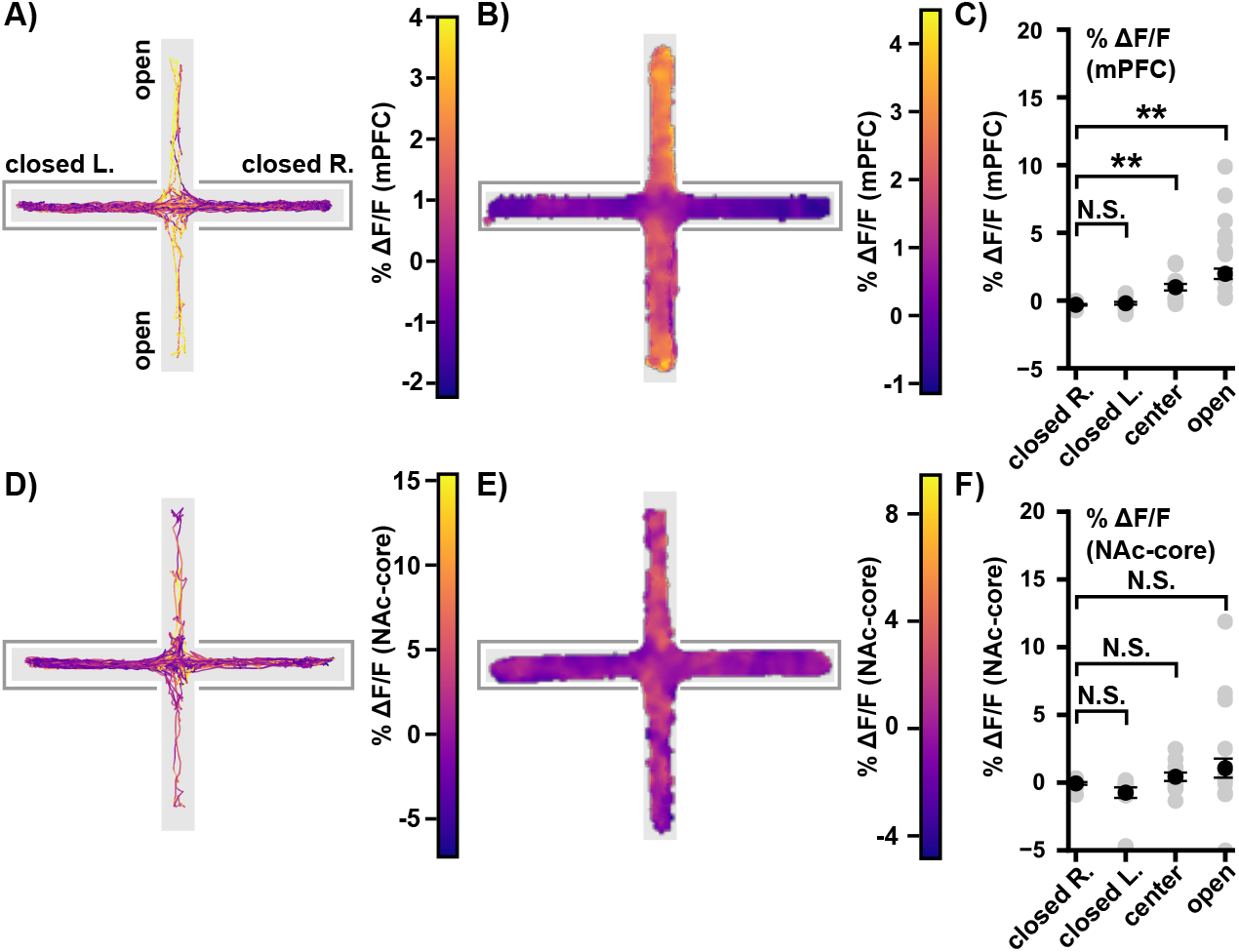
mPFC fluorescence increases as mice explore the open arms, but not a novel closed arm of the elevated plus maze. (* *p <* 0.05, ** *p <* 0.01). **A)** mPFC fluorescence as a single mouse explored the EPM. Line shows the path of the mouse, color the fluorescence. **B)** Gaussian kernel average mPFC fluorescence as a function of position in the EPM. **C)** We found a significant dependence of fluorescence on position (repeated measures ANOVA, *F* = 24.58, *p* = 4.3 *×* 10^−9^). The right arm (closed R.) was where the animal was confined before the experiment began. The left closed arm (closed L.) was the opposite closed arm. Comparing fluorescence in closed R. with the other arms, we found significant increases in the center (pairwise *t*-test with Bonferroni correction multiple comparisons, *t* = 4.93, *p* = 0.0008) and open arms (*t* = 5.53, *p* = 0.0003), but not the opposite closed arm (*t* = 0.83, *p* = 1.0). **D)** Same as panel A, but for recordings in the NAc core. **E)** Same as panel B, but for recordings in the NAc core. **F)** We found a significant dependence of fluorescence on position for the NAc core recordings (repeated measures ANOVA, *F* = 3.39, *p* = 0.03); however, comparing the closed arm where mouse was initially confined with the other three arms, we found no significant differences for the opposite closed arm (pairwise *t*-test with a Bonferroni correction for multiple comparisons, *t* = − 1.51, *p* = 0.48), center (*t* = 1.17, *p* = 0.81) or open arm (*t* = 1.38, *p* = 0.59)

We found significantly higher fluorescence in both the center and open arms of the maze, but did not observe increases in mPFC dopamine as mice entered the closed arm opposite to where they were initially confined (Fig. 3C), suggesting that aversion, but not exploration or novelty, elicit dopamine release in this task.

In recordings in the NAc core, we found a significant dependence of fluorescence on position, though we did not find significance in any pairwise comparisons (Fig. 3D-F). For the open arms in particular, although the mean is elevated, the variance across animals is large with some mice showing decreased fluorescence in the open arm.

Finally, we performed the same analysis for the isosbestic fluorescence, but found no significant dependence of fluorescence on location in either cohort (Fig. S3A-D).

### Dopamine release increases with larger rewards, but is absent at reward omission

To test the responses of mPFC dopamine to simple water rewards, water-restricted mice were placed in a chamber with a single water port at one end and a beam break at the other (Fig. 4A). To get water, mice had to cross the beam break, triggering an auditory cue, and then lick at a water port on the opposite end triggering water release on 50% of trials. We found small, but significant increases in fluorescence following a lick on trials in which the water was delivered and a decrease in fluorescence that did not quite reach significance when no water was delivered (Fig. 4C,D). Although the dip in fluorescence is consistent with a reward prediction error interpretation, we caution that size of this dip is small, less than 0.1% ΔF/F, and a careful examination of the recording shows that it begins slightly before the mouse licks the port. We are confident, however, that there is not an *increase* in dopamine release when the reward is omitted. We observed no increase in fluorescence at the beam break and auditory cue (Fig. 4B). A lack of fluorescence at cues predicting reward has been observed in some studies (6, 7), but not all (3), and may depend on the amount of prior training, which, was only a single day in this task.

**Fig. 4.**
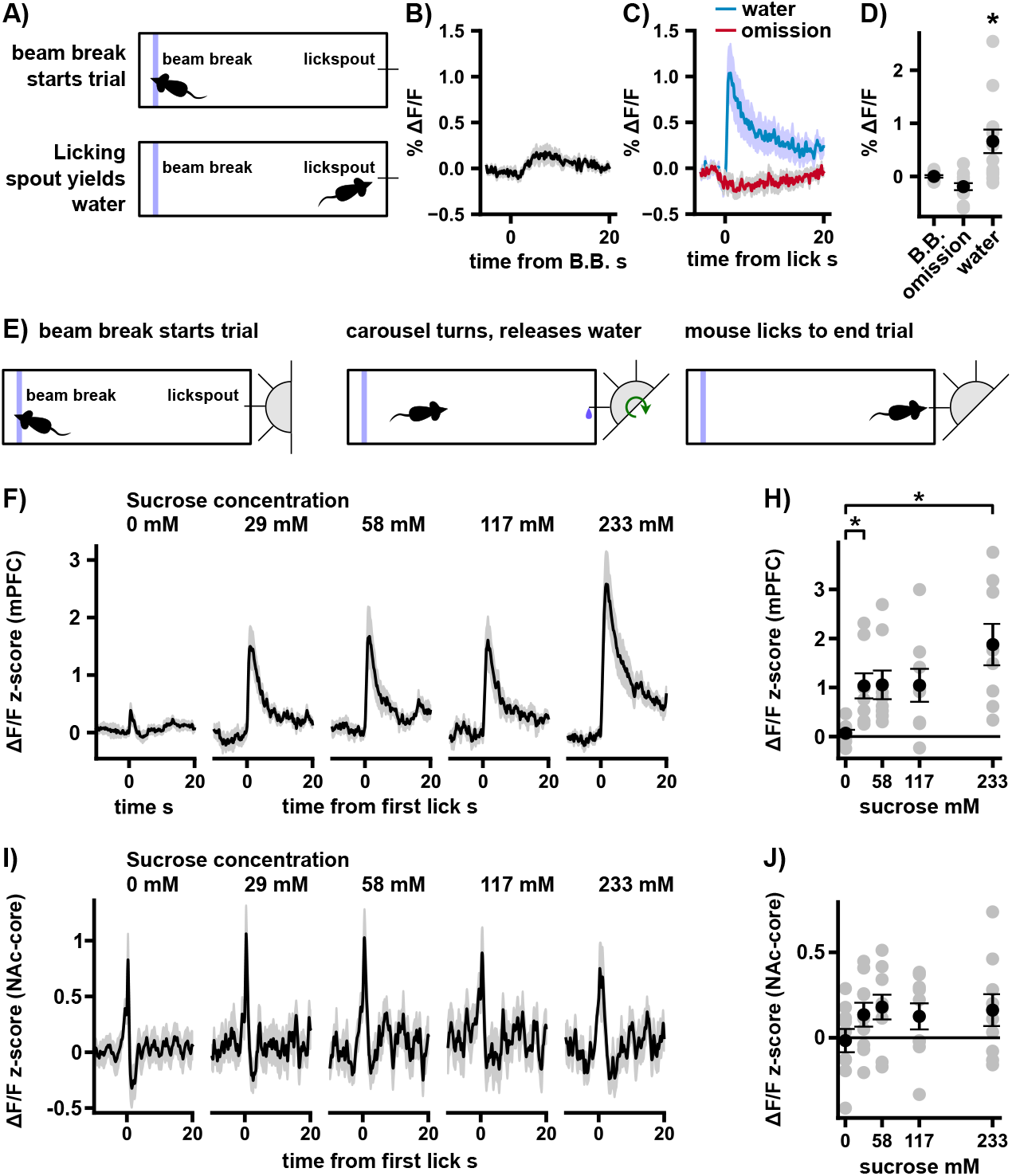
mPFC fluorescence increases following water and sucrose rewards, but not reward omissions. (* *p <* 0.05, ** *p <* 0.01). **A)** Illustration of reward task in which mice crossed a beam break and licked a spout to receive a water droplet. Water was omitted on 50% of trials. **B)** Perievent average mPFC fluorescence around the beam break crossing. Note that the increase in fluorescence is delayed by approximately 2 seconds, after which fluorescence from licking at the port can contribute to the perievent average. **C)** Perievent average mPFC fluorescence at the water delivery or omission when the mouse licked. **D)** The average fluorescence in a 5 second window increased significantly for water (pairwise *t*-test, with a Bonferroni correction for multiple comparisons, *n* = 12, *t* = 2.86, *p* = 0.046), nearly reached significance for a decrease following omission (*t* = −2.78, *p* = 0.052) and was non-significant for the beam-break (*t* = 0.02, *p* = 1.0). **E)** Illustration of a variation of the reward task in which one of five solutions can be delivered from a rotating carousel with five spouts. Mice had to cross a beam-break to trigger movement of the carousel and release of one of the five solutions. Mice had to lick the spout at least once to end the trial. **F)** Perievent average z-scored fluorescence in the mPFC as mice lick the spout receiving either water or one of four concentrations of sucrose. **H)** Summary of mPFC fluorescence averaged over a 5 second window after the first lick. Gray dots represent individual animals. Fluorescence depended significantly on concentration (repeated measures ANOVA, *n* = 8, *F* = 10.36, *p* = 2.8 *×* 10^−5^). Performing all comparisons among water, the lowest and highest concentrations of sucrose, we found that, relative to water, 29 mM sucrose had higher fluorescence (pairwise *t*-test, with a Bonferroni correction for multiple comparisons, *n* = 8, *t* = 3.76, *p* = 0.021) as did 233 mM sucrose (*t* = 4.19, *p* = 0.012), but the comparison of 233 mM sucrose vs. 29 mM sucrose did not reach significance (*t* = 2.84, *p* = 0.075). **I)** Same as panel F, but recorded in the NAc core. **J)** Same as panel H, but for NAc recordings. All concentrations elicited a phasic response at the first lick, but there was no significant dependence of fluorescence on concentration (repeated measures ANOVA, *n* = 9, *F* = 2.24. *p* = 0.09)

To test the effect of increasing the value of the reward, we considered the effect of drinking solutions with either 0 mM, 29 mM, 58 mM, 117 mM or 233 mM sucrose. To be able to deliver an arbitrary solution from this set of concentrations on each trial, we built a rotating carousel with five protruding lick spouts that could be rotated by a servo motor to position one of the spouts at the end of the chamber (Fig. 4E). Trials with sucrose solutions were always interleaved with 1-2 trials of pure water. An example recording from a single mouse is shown in Fig. S4A.

Mice licked significantly more often at ports with higher concentrations of sucrose (Fig. S4B). We found a significant dependence of fluorescence on concentration and significantly more fluorescence at higher concentrations relative to lower concentrations (Fig. 4F,H). We noted that there was considerable variability across trials when the same stimulus was repeated. An example of this variability is shown in Fig. S4E, where the response to the highest concentration of sucrose ranged from around half to twice the average fluorescence over trials. This variation did not necessarily correspond to an overall decrease in fluorescence with repeated trials. We thus observe that, while fluorescence increased with higher concentrations of sucrose, this was only on average, even within animal.

In our recordings in the NAc core, we did not find a significant dependence of fluorescence on concentration (Fig. 5I, J). However, we noted that the temporal pattern of fluorescence for the NAc core recordings was more complex than in the mPFC, with fluorescence ramping up before mice licked at the port, which has been observed in prior studies (29–31), and dipping following the initial lick at the port for trials with water and the lowest concentration of quinine. These effects may not be reflected by the average over 5 seconds of fluorescence following the first lick that we used in our analysis. Comparing the findings in this section with Fig. 1C, we note that the lack of a response on omission of the reward makes it unlikely that model in which all prediction errors lead to release could describe mPFC dopamine release. The one inconsistency between our recordings and the models in Fig. 1C is the lack of a response at the CS, which, in the two experiments performed here, were either a tone or the turning of the carousel, which makes a noise that could be considered a cue (Fig. S5F). However, mice did not have extensive prior experiences in these tasks and it may take longer for a cue-evoked release event to occur. In line with this possibility, we also did not see a phasic rise in fluorescence at the cue in the NAc core recordings (Fig. S5G), though fluorescence does begin ramping up at this point.

**Fig. 5.**
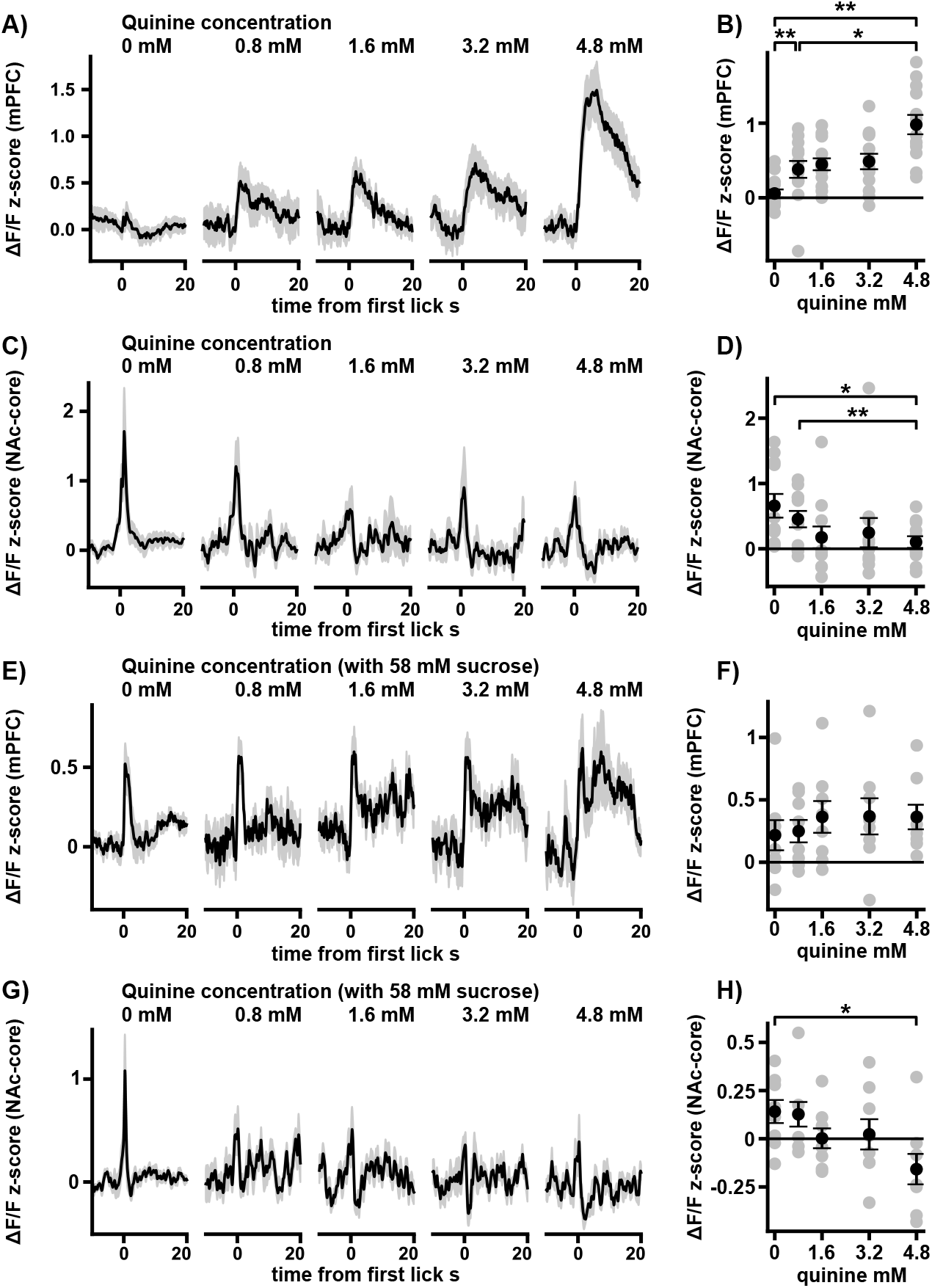
Adding aversion to a rewarding stimulus increases fluorescence. (* *p <* 0.05, ** *p <* 0.01). **A)** Perievent average z-scored mPFC fluorescence for water and four concentrations of quinine. **B)** Summary of average fluorescence over a 5 second window. Gray dots represent individual animals. Average fluorescence depended significantly on concentration (repeated measures ANOVA, *n* = 13, *F* = 10.71, *p* = 2.8 *×* 10^−6^). We performed all pairwise tests between water and the highest and lowest concentrations of water. We found that all were significantly different (pairwise *t*-test with Bonferroni correction for multiple comparisons; water vs. 0.8 mM quinine, *n* = 13, *t* = 3.87, *p* = 0.0067; water vs 4.8 mM quinine, *t* = 5.85, *p* = 0.00023; 0.8 mM quinine vs 4.8 mM quinine, *t* = 3.38, *p* = 0.016). **C)** Same as panel C for NAc core. **D**) Same as panel D for NAc core. Average fluorescence depended significantly on concentration (repeated measures ANOVA, *n* = 11, *F* = 2.72, *p* = 0.043). Both water and the lowest concentration of quinine had significantly higher average fluorescence than 4.8 mM quinine (pairwise *t*-test with Bonferroni correction for multiple com-n parisons; water vs. 4.8 mM quinine, *t* = 3.07, *p* = 0.035; 0.8 mM quinine vs. 4.8 mM quinine *t* = 4.56, *p* = 0.0031), but water was not significantly different from 0.8 mM quinine (*t* = 1.48, *p* = 0.51). E) Perievent average response to various quinine concentrations in the presence of a constant concentration of 58 mM sucrose). **F)** Summary of average fluorescence over a 5 second window. We found no significant dependence of fluorescence on concentration (repeated measures ANOVA, *n* = 8, *F* = 0.59, *p* = 0.67). We noted that, while the initial phasic response to the five solutions was similar, the fluorescence appears elevated after this time. This feature is discussed in the text and in Fig. S5C **G)** Same as panel G for NAc core. **H)** Same as panel H for NAc core. Average fluorescence had a significant dependence on concentration (repeated measures ANOVA, *n* = 8, *F* = 4.22, *p* = 0.0084). Pure sucrose elicited a significantly larger response than sucrose mixed with 4.8 mM quinine (pairwise *t*-test with Bonferroni correction for multiple comparisons, *t* = 4.28, *p* = 0.04.) while the other two comparisons were not significant (pure sucrose vs. sucrose and 0.8 mM quinine, *t* = 0.24, *p* = 1.0, 0.8 mM quinine vs. 4.8 mM quinine, *t* = 2.57, *p* = 0.11)

### Adding aversion to a rewarding stimulus increases release

We next tested the effect of mixing reward and aversion in a single stimulus, which we will refer to as a mixed valence stimulus. We sought a pair of stimuli, one rewarding and one aversive, which could be simultaneously delivered to an animal without the rewarding and aversive components masking each other.

We noted that animals in the natural world must seek water to survive, but water sources can be contaminated with toxins, forcing animals to carefully weigh their thirst needs against potential harmful effects. In such scenarios, the taste of the solutes in the water may be unpleasant, but this taste does not impair the ability of the mouse to detect that it is consuming water.

Following this line of reasoning, we sought a collection of stimuli that started with a purely rewarding stimulus (water) to which we could add an increasingly large concentration of an aversive solute. We note that if mPFC dopamine is a function only of net valence (reward - aversion), dopamine release must be zero at a stimulus somewhere between the purely rewarding and highly aversive stimulus (Fig. 1A, green). However, if it has a more general dependence on reward and aversion, such as total valence (reward + aversion), it may have no dip between the two extremes (Fig. 1A, orange).

Using the carousel task described above, we recorded the responses of the mPFC dopamine system in water-restricted mice as they drank water mixed with various concentrations of the highly bitter substance quinine (32). Our solutions ranged from pure water and a slightly bitter solution (0.8 mM quinine) to an intensely bitter solution (4.8 mM quinine). The perceived quality of the different solutions was measured by number of licks that mice made at the spout for each solution, which decreased significantly at higher concentrations of quinine (Fig. S5A).

An example recording is shown in (Fig. S5C). As the concentration of quinine was increased from pure water to the solubility of quinine, we observed a significant increase in fluorescence that was highest for the highest concentration (Fig. 5A, B). We note that there was a significant increase in fluorescence even from water to the lowest concentration of quinine. We compared these recordings with recordings in the NAc core region and observed, as expected, that fluorescence decreased at higher concentrations of quinine (Fig. 5C, D). We note that, at the lowest concentration of quinine, there was still a phasic increase in fluorescence following consumption of the solution and the average fluorescence was significantly higher than for the highest concentration of quinine.

This, together with the elevated lick rate for the lowest concentration of quinine relative to the highest, suggested to us that this stimulus has a net positive valence, even if it is a smaller net valence than pure water. The increase in dopamine-dependent fluorescence from water to the lowest concentration of quinine is thus a challenge to any theory that supposes that mPFC dopamine release only depends on the net valence of a stimulus.

We also considered the effect of adding sucrose to all of the solutions, while maintaining the same five concentrations of quinine. We note that this experiment is not as clean as the previous experiment as the sweetness of the solution may mask the taste of the quinine and vice versa.

We found that, as in the previous experiment, as the concentration of quinine is increased, the number of licks at the spout significantly decreased (Fig. S5B). As in the pure quinine experiment, we found no dip in the fluorescence as the concentration of quinine was increased (Fig. 5E, F). We noted that the time-dependence of the fluorescence signal appeared to have two phases in this experiment, with an initial peak present at all concentrations of quinine, and presumably due to the background sucrose, followed by a large plateau of increased fluorescence that was present at higher concentrations of quinine. We found no significant dependence of fluorescence averaged over the first five seconds after the first lick at the port (Fig. 5F). As a post hoc analysis, we also examined a window from 5-10 seconds to try to capture the increasingly large plateau and found a significant dependence of fluorescence on concentration (Fig. S5C).

Repeating this experiment in the NAc core, we found that, as expected, increasing the concentration of quinine significantly decreased fluorescence (Fig. 5G,H), consistent with NAc core dopamine being a function of net valence.

We again observed a high amount of variance in the mPFC fluorescence responses to quinine, as we did for the pure sucrose experiments, with the fluorescence varying by around a factor of two and a complex dependence on trial number (Fig. S5F). As in the pure sucrose experiments, our claims of an increase in fluorescence with higher concentrations of quinine should thus only be taken as a statement of the average, not individual trials.

Comparing our mPFC recordings to Fig. 1C, purple region, we see that an increasing response to the addition of aversion to a rewarding stimulus is consistent with a total valence model, but difficult to reconcile with any model that is a function of net valence alone. We note that the NAc core recordings, however, are consistent with RPE, which depends only on net valence; the more aversion is added to the rewarding stimulus, the lower the fluorecsence.

Finally, we repeated the analysis in this section for the isosbestic fluorescence recorded in the mPFC and NAc core cohorts and found no significant dependence on concentration in either the first quinine experiment or the sucrose and quinine experiment (Fig. S5D,E).

### Release following novel object and social interactions is better explained by valence than engagement

In the previous experiments we saw that adding an aversive stimulus to a rewarding stimulus did not decrease fluorescence as expected if dopamine depends entirely on net valence. These experiments suggested the possibility that mPFC dopamine could depend primarily on how salient the stimulus was, rather than on valence. We thus measured the change in dopamine-dependent fluorescence following an array of stimuli consisting of novel objects and social interactions, some of which had a high valence, while others had no obvious valence, but were nonetheless engaging to the mice, as measured by the amount of time they spent interacting with the stimulus (Fig. 6F).

**Fig. 6.**
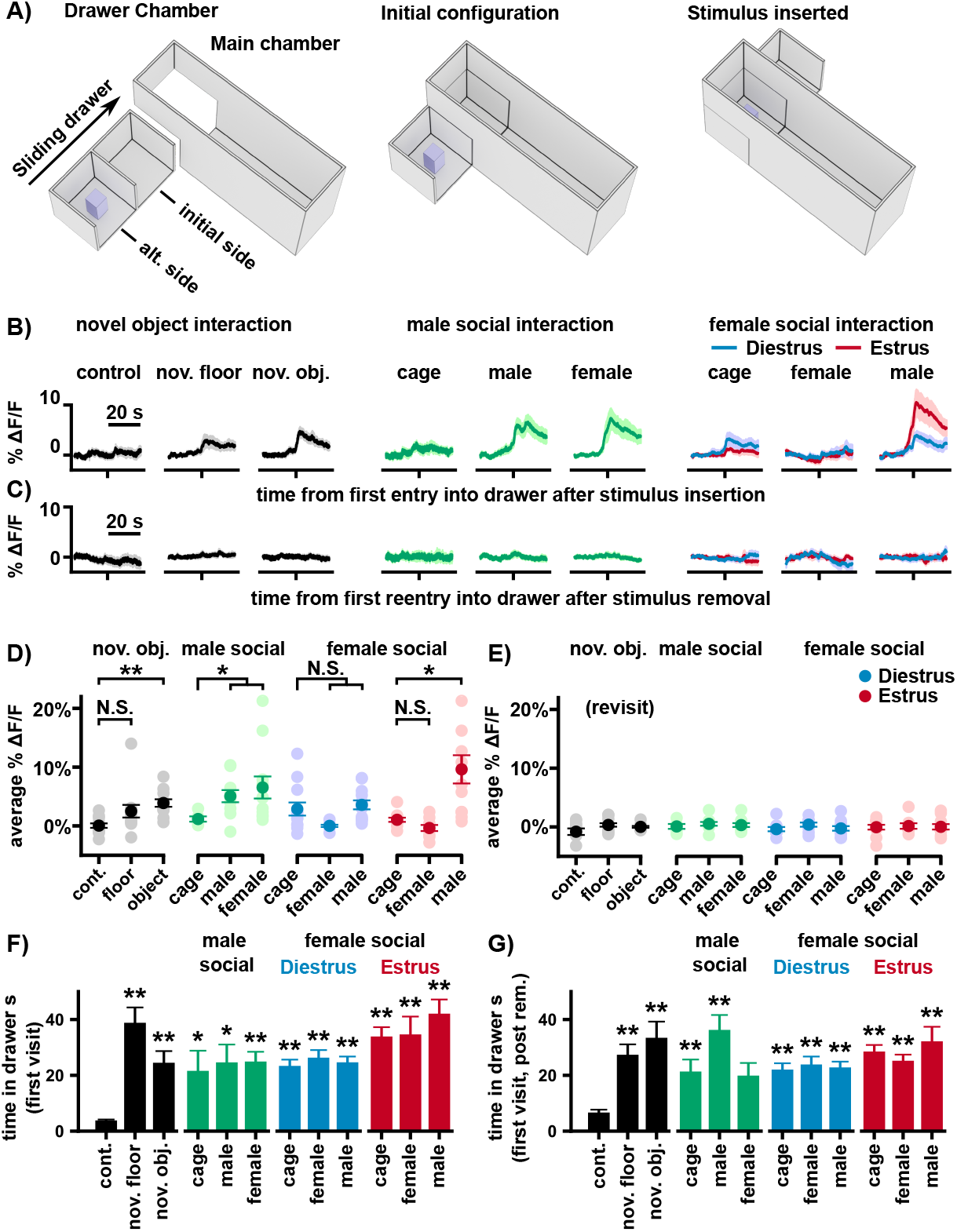
Valence and not engagement are linked with dopamine release. (* *p <* 0.05, ** *p <* 0.01). **A)** Illustration of the drawer chamber used in the task allowing surreptitious insertion of stimuli into a chamber with a mouse. Experiments were run in red light and drawer was only slid when the mouse was at the opposite end of the chamber. **B)** Perievent average fluorescence around the first entry intro the drawer after stimulus insertion. The control was a swap of the drawer with no stimulus. For social experiments, mice were previously exposed to the empty cage to reduce any signals from novelty. Female mice in diestrus and estrus are shown in blue and red respectively. Note the especially high fluorescence for a female mouse in estrus meeting a male mouse. **C)** Same as B but for the first entry after the stimulus was removed. Note the lack of response. **D)** Average fluorescence over the first 5 seconds after drawer entry. Light colored dots are individual animals. Stimuli were analyzed in four groups, non-social, male social interaction and female social interaction in estrus and diestrus. We found that all groups had a significant dependence on condition (one-way ANOVAs, non-social *n* = 11-12, *F* = 16.34,, *p* = 0.00028; male-interaction, *n* = 5-11, *F* = 11.24, *p* = 0.0036; female interaction (diestrus), *n* = 9-12, *F* = 15.47, *p* = 0.0043; female interaction (estrus), *n* = 9-11, *F* = 11.42, *p* = 0.003). Fluorescence for stimuli were compared with a control stimulus consisting of either a drawer swap with nothing in it for non-social stimuli or an empty cage for social stimuli. We found significance increases in fluorescence for the novel object (independent *t*-test, *t* = 4.94, *p* = 0.00027), a male mouse meeting either a male or female mouse (*t* = 3.24 or 2.64, *p* = 0.014 or 0.045), and a female mouse meeting meeting a male mouse in estrus (*t* = 3.37, *p* = 0.013). **E)** Same as panel D for the average over the first 5 seconds following reentry after stimulus removal. We found no significant dependence of fluoresence on groups (one-way ANOVAs, non-social, *F* = 2.77, *p* = 0.25; Male interaction, *F* = 0.53, *p* = 0.77; Female interaction in diestrus, *F* = 2.05, *p* = 0.36; Female interaction in estrus, *F* = 0.08, *p* = 0.95). **F)** Interaction times with the stimulus. Interaction times were compared with the control drawer swap (independent *t*-test, corrected for multiple comparisons). **G)** Interaction times during the first revisit to the drawer after stimulus removal. Note that all but one stimulus had stimulus interaction times significantly larger than the control swap (independent *t*-test, corrected for multiple comparisons).

In order to introduce stimuli into a chamber without a mouse being aware of the insertion, we designed a box in which one end had a “drawer” that could swap the end of the chamber with an copy (Fig. 6A). Recordings were aligned at the moment when mice first entered the drawer after insertion of a stimulus or after its removal.

We began by comparing changes in fluorescence evoked by inserting nothing (a control stimulus), a novel floor made of the same material as the chamber but with a different texture, or a novel object. We found a significant dependence on stimulus type and a significant increase in fluorescence for the novel object, but not the novel floor (Fig. 6B, D). We noted that mice spent significantly longer investigating the novel floor than a control stimulus, the second longest investigation time of all of the stimuli we inserted (Fig. 6F), suggesting that the floor was engaging despite the dopamine response being the smallest of all the non-control stimuli.

We then tested the effect of social interactions on mPFC dopamine release. We found a significant dependence on stimulus between a male mouse investigating an empty cage which mice had previously been exposed to and a cage with a male or female mouse, with both male and female mice producing significantly higher fluorescence than the empty cage (Fig. 6B, D).

For social experiments in female mice, prior work has shown that sexual attraction depends on the estrus cycle (33–35). We tracked the estrus cycle of female subjects using stained vaginal smears (Fig. S6C) (36) and performed recordings during either estrus, when female mice are most receptive to mating, or diestrus. In both conditions we found a significant dependence of fluorescence on stimulus and for mice in estrus a significantly higher fluorescence than the empty cage control when a female mouse met a male mouse. This particular fluorescence was the highest fluorescence we recorded across any of our reward experiments and second only to the response to the US in our fear conditioning experiment.

Examining the investigation times for each stimulus, we found that mice spent significantly more time investigating every stimulus, when compared with the control stimulus consisting of an empty drawer (Fig. 6F). We noted that the highest average fluorescences observed for the male and female mice were for interacting with mice of the opposite gender, both highly appetitive stimuli, while stimuli that did not have an obvious valence, like a female mouse interacting with a female mouse or mice of either gender interacting with a novel floor led to the smallest increases in dopamine.

We also measured the effect on fluorescence of a mouse discovering that a stimulus had been removed from the chamber. Examining the average fluorescence over the first 5 seconds after entering the drawer post stimulus removal, we found no significant dependence of fluorescence on stimulus across any of the conditions (Fig. 6C, E). This was not due to mice not realizing that the stimulus had been removed, as there was a significant increase in investigation time of the drawer post removal for all but one of the stimuli relative to the control stimulus (Fig. 5G). We thus find that, as with painful stimuli and water rewards, omission of a stimulus did not elicit an increase in fluorescence in our experiments.

Finally, we repeated our analysis for the isosbestic fluorescence and found no significant dependence of fluorescence on condition (Fig. S6A,B)

## Discussion

Our experiments are consistent with the following observations about the mPFC dopamine system: **1)** As previously reported, dopamine is released for both rewarding and aversive stimuli, with painful stimuli releasing particularly large amounts of dopamine. We note, however, that outside of painful stimuli, the largest increase in fluorescence observed was from a highly rewarding stimulus, a female mouse interacting with a male mouse while in estrus. **2)** When rewarding and aversive stimuli are combined, the fluorescence is inconsistent with any function of the net valence of the stimulus and is qualitatively similar to a sum of rewarding and aversive components. **3)** Omissions of expected stimuli do not produce dopamine release. We note that this finding is a challenge to theories of mPFC dopamine that propose that dopamine alerts the mPFC to changes in the environment or that claim that dopamine is released when stimuli are interesting, salient or novel. **4)** Novel stimuli and environments do not elicit dopamine release in the absence of valence, even when the stimuli are engaging. One possible exception to this rule is the novel object experiment, which, unlike the novel floor experiment, did significantly increase fluorescence. However, mice typically interact cautiously with novel objects suggesting that this stimulus may be aversive because of neophobia, which we do not observe as mice explore a novel floor.

Based on these findings, we propose that stimulus-elicited mPFC dopamine release is a function of the total valence of the stimulus

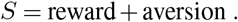

Examining Fig. 1C, we see the principle advantage of this model over other models that we considered is that *S* is positive for both reward and aversion and reward and aversion do not cancel. For unconditioned stimuli, it agrees with all of our recordings.

Total valence is the sum of two signals, one for reward and one for aversion. A simple mechanism for producing such a signal would be to have two populations of dopamine neurons whose axons project to the mPFC, one of which releases dopamine in proportion to how rewarding a stimulus is and the other in proportion to how aversive it is. In line with this, a recent study suggested that dopamine neurons that project to the mPFC are biased to respond to either reward or aversion (6). Although this finding does not imply our proposal, as the reward-biased and aversion-biased populations may inhibit each other, it would imply it if the two populations are largely independent.

### Total valence and prediction errors

If mPFC dopamine responds to the total valence of a stimulus, it is natural to guess, based on the reward prediction error (RPE) theory of mesolimbic dopamine, that mPFC dopamine release should correspond to a similar theory, which we will call *S*PE, that depends on the total valence, *S*, instead of *R*. Explicitly, we define, in discretized time, the temporally-discounted sum of expected future total valence,

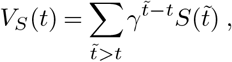

where *S*(*t*) is the predicted total valence of stimuli received at time *t* and *γ* is the temporal discounting factor. We then define, in analogy with the formula for RPE (11) from temporal difference learning (37, 38),

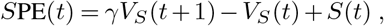

and hypothesize that mPFC dopamine neuron activity corresponds approximately to *S*PE(t). Predictions of this model were shown in Fig. 1C. Here we discuss evidence for and against *S*PE.

Cues predicting rewarding events can elicit dopamine release in the mPFC (3); however, this finding is not consistent across the literature, with some studies finding association-driven release primarily for aversive stimuli and not rewarding stimuli (6–8). Indeed, in this study, there is a clear increase in association-driven dopamine release in cued fear conditioning, but not for a cue predicting a water reward. A major difference between the reward task described here and the task in (3) is the length of training, with only a single day in this study and more than a week in the prior study. This leaves open the possibility that prediction errors in the mPFC dopamine system may form more quickly for aversive stimuli than rewarding stimuli. Supporting this hypothesis, in our NAc core recordings, a phasic increase in fluorescence at a cue predicting sucrose-water had not yet formed in our carousel experiment (Fig. S5G), but more research is needed. One finding in this study that agrees with *S*PE is the lack of responses on omission of expected stimuli. Because *S*(*t*) is always positive, any prediction error on omission must be negative. Given the slow kinetics of mPFC dopamine, a short pause in the firing of mPFC dopamine neurons would likely be undetectable. Voltage imaging or electrophysiological recordings from mPFC-projecting dopamine neurons could confirm whether there are pauses in firing or whether omissions have little effect on both dopamine concentrations and dopamine neuron spiking, which would be inconsistent with *S*PE.

Another aspect of our recordings that challenges a simple model like *S*PE is the high variability of responses we observed in our recordings. Examining the release of mPFC dopamine upon licking pure water, or water combined with solutes that are either rewarding or aversive, we often find doubling or halving of the fluorescence signal from trial to trial relative to the average. This suggests to us that other latent variables are at play which influence dopamine release. One recent study suggested that repeated stimuli rapidly decrease the amount of mPFC dopamine release (8), but we do not consistently observe a decrease here, perhaps because, in most of our experiments, trials with the same stimulus are interleaved with different stimuli, reducing habituation and maintaining mPFC activity related to the task. Nonetheless, there may be a strong attentional component to mPFC dopamine release that diminishes responses to familiar cues, even if they occur at unpredictable times.

The variability we observe may also arise from internal states of the mPFC, which sends excitatory projections back to mPFC-projecting dopamine neurons exciting them (39), or to local interneurons that inhibit dopamine neuron activity. This top-down modulation could be affected by increased or decreased frontal attention to the stimulus or may arise from cognitive variables that we are not measuring or considering in our experiments.

Despite these challenges, examining Fig. 1C, we find that *S*PE makes qualitatively similar predictions to our recordings for all but one of the stimulus types considered. Thus, while we expect our model will be refined in future experiments, we believe it is a strong foundation for building a complete theory of mPFC dopamine release.

### The role of total valence in the mPFC

We close with a brief discussion of why total valence might be a useful signal for the mPFC. Total valence increases whenever the amount of available rewards or the total danger in an environment changes. It differs from a surprise signal in that it does not increase when rewarding or aversive stimuli are omitted, only when the total amount of reward or aversion increases. It also does not respond when stimuli are neutral, having neither a rewarding nor aversive component. These non-responses suggest times when mPFC dopamine does not play an important role in shaping behavior, which may refine existing theories of its role in cognition.

The mPFC dopamine system has been tightly linked with cognitive flexibility (1, 3, 40–47). A positive change in total valence may represent a signal to the mPFC that rewarding or aversive stimuli have appeared in an environment, triggering flexible behavior to adapt to their presence. The opposite of this, when rewards or threats disappear from an environment, might also trigger flexible behavior, but our recordings suggest that this is likely not mPFC dopamine dependent, a prediction that could be tested in future studies.

From this prediction, it would follow that mPFC dopamine may be more important for loading new goals and threats into working memory as they appear in the environment, rather than removing them when they are no longer relevant. This idea is consistent with a curious result in (3) in which it was found that mPFC dopamine stimulation was effective at inducing explorative behavior when paired with choices that were unrewarded, causing mice to select them more often, but did not disrupt behavior when paired with correct choices. When paired with incorrect choices, the stimulation may have increased the representation of alternative strategies that select the unrewarded choice. When mPFC dopamine release was paired with correct choices, it did not have an effect, perhaps because mPFC dopamine release did not remove or disrupt a strategy already established in working memory.

In summary, our recordings suggest that mPFC dopamine release can be interpreted as a kind of surprise signal, but one restricted to a narrow class of surprises, the unexpected addition of new rewards or threats in an environment. Future studies can test if this total valence model holds when extended to behavioral tasks that require cognition, such as the storage of an item in short-term memory (48, 49) or a switch from one strategy to another, both of which are believed to be mPFC dopamine dependent.

## Methods

All experiments were conducted in accordance with procedures approved by the Institutional Animal Care and Use Committee (IACUC) at Cornell University. All mice used in the study were C57BL/6.

### Injections and implantations

To drive expression of GRAB_DA_3h or GRAB_DA_3m, mice were sterotactically injected with 500 nL of 1013 vg/mL AAV9 at 150 mL/min in either the mPFC or NAc core (pAAV-hsyn-rDA3h or pAAV-hsyn-rDA3m, WZ Biosciences) using methods described previously (3). Stereotactic coordinates for the mPFC were (1.75 A.P., 0.5 M.L., −2.55 D.V.) and for the NAc core (1.0m AP, 1.0 MD, −4.2 DV). We note that taking the average of the coordinates measured after histology gives a mean for mPFC of (1.65 AP, 0.47 ML, −2.75 DV), which is around 200 *µ*m ventral to the target coordinates, and (1.16 AP, 0.91 MD, 4.21 DV) for NAc core. After injection, mice were implanted with 400 *µ*m diameter low fluorescence optical fibers (Doric Lenses) 200 *µ*m above the injection site. No experiments were performed until at least 4 weeks post-surgery to allow for expression.

### Fiber photometry recordings

To record fluorescence, we used a custom-built fiber photometry rig based on previous designs (18–20). Excitation and Isosbestic light were generated by 470 nm and 415 nm LEDs (Thorlabs) respectively, passing through 488 nm and 420 nm bandpass filters (Semrock). The two light sources were joined with a 425 nm dichroic mirror (Thorlabs) before being reflected off a 500 nm dichroic mirror onto a collimating lens which focused the light on a 400 *µ*m diameter fiber optic patch cable that was attached to the implant of the experimental subject. Emitted light was passed through a 525 nm step filter (Semrock) and focused onto a femtowatt silicon photoreceiver (Newport). The excitation and isosbestic light sources were sinusoidally modulated and lock-in amplification was performed with an RZ5D base processor (Tucker-Davis Technologies).

### Fear conditioning

Fear conditioning was performed in a custom built chamber with a metal shock floor (MedAssociates). Mice were placed in the chamber 5 minutes before photometry recordings began and an additional 5 minutes elapsed before the task started. The conditioned stimulus (CS) consisted of either a 5 second 10 kHz tone or 5 seconds of white noise, while the non-conditioned stimulus (NCS) consisted of the other option. During conditioning, the first eight CS presentations were terminated by a 500 ms 0.5 mA footshock. Each presentation of the CS was paired with a presentation of the NCS, with the order of the stimuli decided randomly for each pair. The inter-stimulus interval was drawn randomly from a uniform distribution between 2 and 2.5 min. The extinction phase, which immediately followed the conditioning phase, consisted of 30 presentations of the CS without the US using the same inter-stimulus interval. On the following day, this same extinction protocol was repeated as a test of extinction memory.

### Elevated plus maze

The elevated plus maze (EPM) consisted of a custom-built chamber with two open arms and two closed arms surrounded by high walls. The mouse was placed in one of the closed arms initially with a divider blocking the entrance to the arm. During the experiment, mice were initially isolated in the closed arm for the first 5 minutes before the divider was removed, allowing the mouse to explore the maze for the next 25 minutes. The maze was illuminated by 10 W white light and surrounded by white curtains. Behavior was filmed with a camera mounted above the maze and mice were tracked with ANY-maze.

### Carousel Chamber

To deliver up to five different solutions to mice, we built a rotating carousel of lick ports. The rotation system consisted of a small servo motor mounted at the base of a vertical metal shaft supported by bearings. Mounted on the shaft were 5 radially projecting spouts that received water from five 15 mL centrifuge tubes. Flow to the spouts was controlled by five solonoid valves. Licks at the spouts were recorded with a custom capacitative lick sensor attached to all five ports. To disguise which port was being selected on each trial, we balanced the amount of time that the carousel turned so that it was the same for any of the switches. Hence, the carousel rotated past its intended target and rotated back or, on trials when the port was the same as on the previous trial, the carousel rotated away from the port and then back. The movement of the carousel took approximately 800 ms. The lowest concentration solution (either water or pure sucrose) was always attached to the middle of the five ports and in all our experiments 1-2 trials with the lowest concentration interleaved any trials with higher concentrations. To complete a trial, mice had to cross a beam break at the opposite end of the chamber and then lick at the port. Mice were trained in the task with only water on multiple days until they could achieve 60 trials over a 30 minute period.

### Drawer Chamber

To surreptitiously insert stimuli into a chamber holding a mouse, we developed a chamber with a drawer in the back that could be slid to swap the back of the chamber with an identical copy. The chamber was built from PVC that was smooth on one side, which was used for the interior of the chamber and drawer, and had a faux wood grain texture on the other. The bottom of the drawer was slightly lower than the floor of the chamber and two squares of PVC were placed into the drawer for all experiments so that, with the insert, the floor of the chamber was level with the drawer. For the novel floor experiment, one of these inserts was flipped upside down, revealing the wood grain texture. A control swap consisted of moving the duplicate of the drawer into the chamber with nothing in the drawer and the same floor surface as the chamber. The novel object was a small connector used for mounting optical parts (Thorlabs) and the cage was a small pencil holder. Different cages were used for the empty cage and male and female social interaction experiments. A small weight was placed on the cage to hold the cage still. Mice used in the social experiments were selected to be of similar age (within 2 weeks), but not litter mates. All experiments were conducted in dim red light. Mice were placed in the chamber for 30 minutes on the first day and then for 30 minutes with the empty cage for the next two days to habituate them to the chamber and empty cage. During recording, the mouse was in the cage for 15 minutes before the drawer was swapped. The stimulus remained in the cage for 10 minutes before being removed. The drawer was only swapped when the mouse was at the opposite end of the chamber. For experiments in which a female mouse was interacting with either a empty cage or another mouse, the empty cage control and social interaction were performed as successive swaps, with the order randomized in order to ensure that both were recorded in the same phase of the estrus cycle. Interaction times were measured as the amount of time mice spent in the drawer region before leaving. Two datapoints were removed as outliers from the interaction times using Grubbs’s test, one from a mouse that spent 5 minutes in the drawer with a novel object and the second from a mouse that spent two minutes in the drawer after removal of the novel floor.

### Estrus cycle tracking

Tracking was performed using stained vaginal smears (36). Vaginal smears were examined under a light microscope and estrus and diestrus were distinguished by the presence of anucleated epithelial cells vs. nucleated cells and neutrophils. Two days before tracking was started, dirty bedding from male cages was introduced into the cages of female mice to ensure cycling. The tracking was started 7 days before photometry recordings to keep consistency and accuracy for prediction. The experiment was performed on a predicted estrus/diestrus day, and the estrous stage was confirmed after the experiment.

## ACKNOWLEDGEMENTS

We would like to thank Andrew Bass and Melissa Warden for comments on a draft of the manuscript. This work was supported by a BBRF young investigator award 139526.

## DATA AVAILABILITY

Preprocessed data as well as the scripts used to produce the figures in the paper are available from a GitHub Repository https://github.com/iellwood/PFCDAAndTotalValence/. To reduce the size of the files, the original recordings at 1000 Hz have been filtered and downsampled to 100 Hz, but are available upon request.

## DECLARATION OF INTERESTS

The authors declares no competing interests.

## Supplementary Figures

**Fig. S2.**
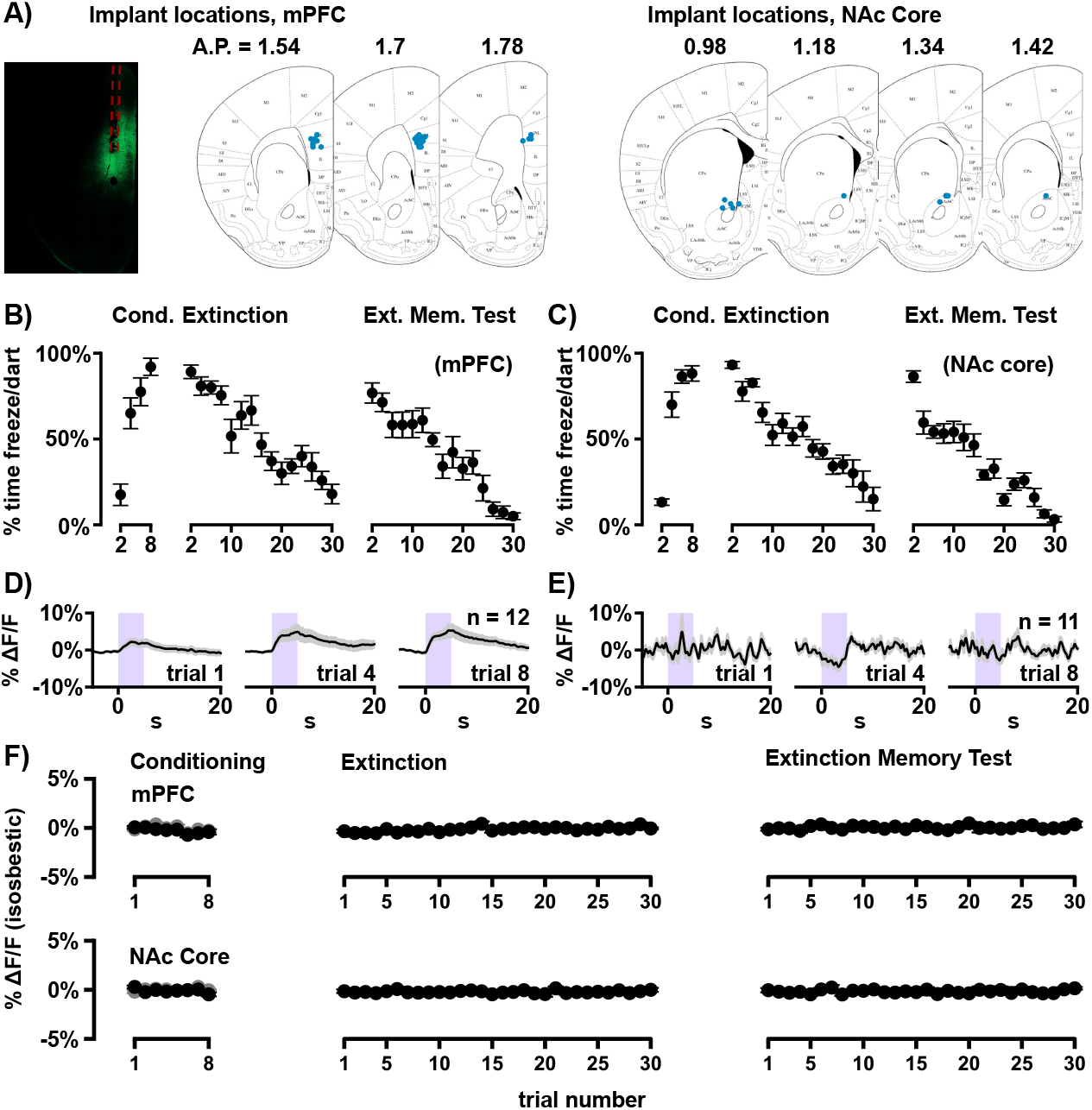
Implant targeting and additional figures from the fear conditioning experiment. (* *p <* 0.05, ** *p <* 0.01). **A)** Image of an example implant (estimated 400 *µ*m diameter implant shown in red) and GRABDA expression (green). Implant locations superimposed on images from Paxinos and Franklin 2001. Note that, while a majority of implants were located in the prelimbic cortex, some were in the dorsal part of IL. Additionally, our photometry rig likely collected fluorescence from both IL and PL. All implant locations in mPFC were in deep layers where the largest number of dopamine fibers reside. **B)** Freezing responses of mice implanted in the mPFC during the fear conditioning experiment. Darting during the CS was scored as 5 seconds of freezing. **C)** Same as panel A for mice implanted in NAc core. **E)** Perievent average fluorescence in mPFC around the non-conditioned stimulus. **D)** Same as panel D for the NAc core. **F)** Average of the isosbestic signal for the fear conditioning experiment. We found significance for one comparison in the isosbestic signal. There is a small, 0.46% Δ*F/F* rise in the mPFC isosbestic signal during the extinction phase (pairwise *t*-test between the average of the first and last 5 trials of extinction, *n* = 11, *t* = 2.26, *p* = 0.047). We expect this change is due to a decrease in freezing and darting movements during the CS. We found no difference between the CS and NCS at the end of conditioning (*t* = 0.5, *p* = 0.63), or the average of the first and last five trials for the test of extinction memory (*t* = 1.14, *p* = 0.28). We found no significant effects for any of the comparisons in the NAc core recordings, including the CS vs. NCS at the end of conditioning (*n* = 11, *t* = 1.74, *p* = 0.11), the first and last 5 trials averaged during extinction (*t* = 0.59, *p* = 0.57) or extinction memory (*t* = 0.79, *p* = 0.45).

**Fig. S3.**
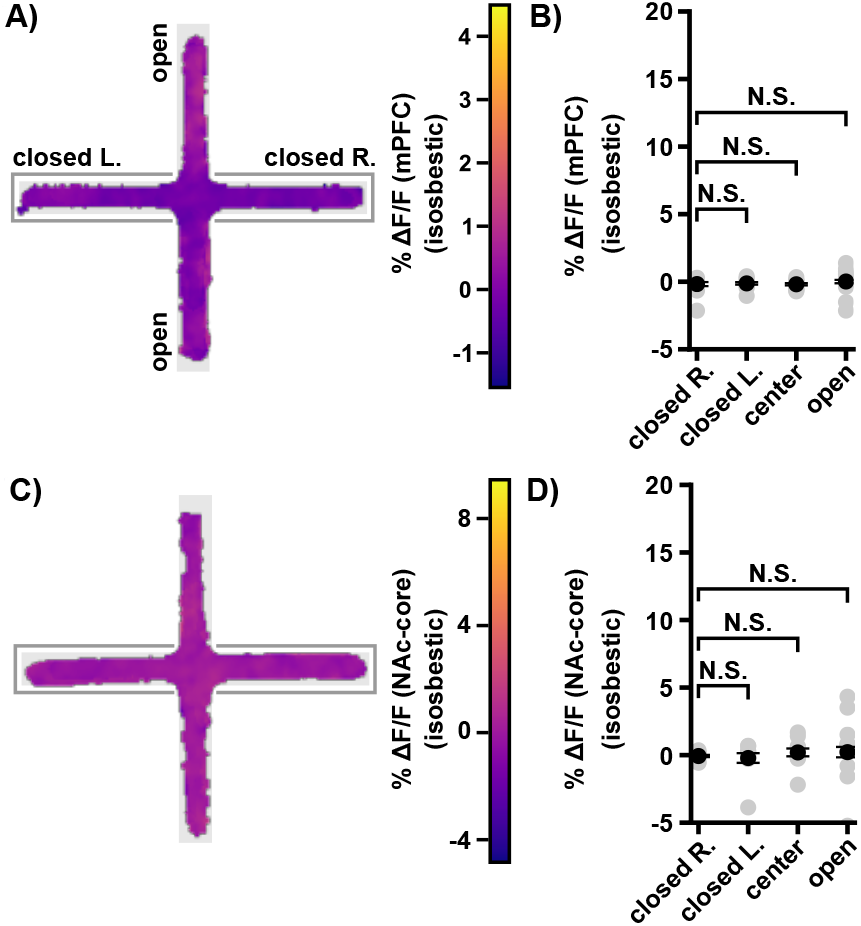
Isosbestic recordings in the EPM. (* *p <* 0.05, ** *p <* 0.01). **A)** Gaussian kernel average of isosbestic fluorescence as a function of position in the EPM. **B)** Average isosbestic fluorescence in four regions of the EPM. We found no significant difference between any of the regions and the right closed arm where the animal was initially confined before the experiment began (pairwise *t* test, with a Bonferroni correction for multiple comparisons; closed L., *n* = 14, *t* = 0.078, *p* = 1.0, center, *t* = 0.078, *p* = 1.0; open, *t* = 0.88, *p* = 0.39). **C)** Same as panel A for isosbestic fluorescence recorded in the NAc core. **D)** Same as panel B for isosbestic fluorescence recorded in the NAc core. We found no significant comparisons (pairwise *t* test, with a Bonferroni correction for multiple comparisons; closed L., *n* = 10, *t* = 0.34, *p* = 1.0, center, *t* = 0.72, *p* = 1.0; open, *t* = 0.61, *p* = 0.39).

**Fig. S4.**
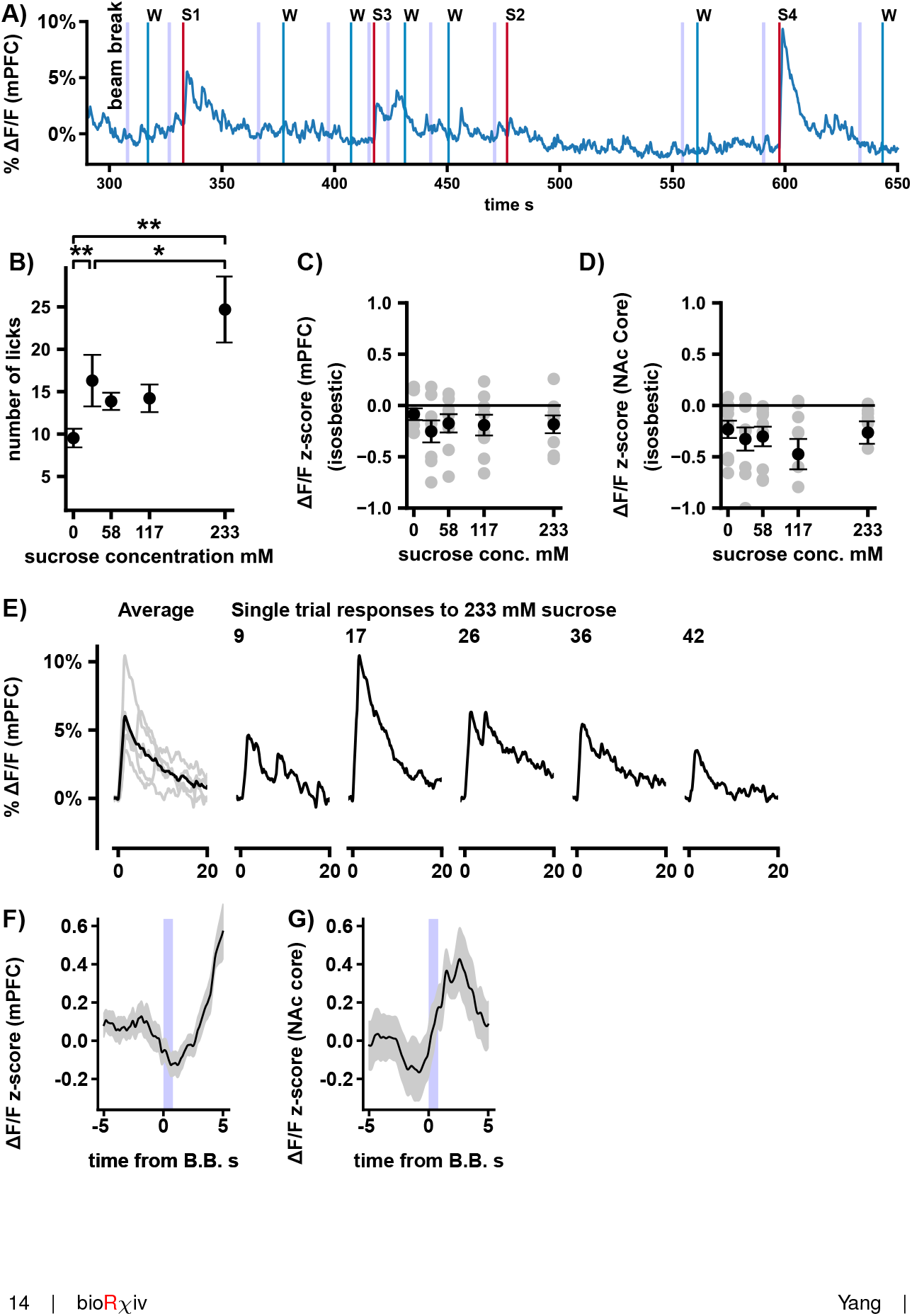
Additional details of recordings during sucrose and water consumption. (* *p <* 0.05, ** *p <* 0.01). **A)** Example recording of a single animal during the water and sucrose task. Vertical lines represent beam break crossings, water and sucrose delivery as indicated. **B)** Average number of licks at ports with different concentrations of sucrose. Lick rate depended significantly on concentration (repeated measures ANOVA, *n* = 17, *F* = 9.05, *p* = 7.4 *×* 10^−6^). Comparing water and the highest and lowest concentrations of sucrose, we found that all three averages were significantly different (pairwise wilcoxon sign rank test with Bonferonni correction for multiple comparisons; water vs 29 mM sucrose, *W* = 47, *p* = 0.0067; water vs. 233 mM sucrose, *W* = 41, *p* = 0.0032; 29 vs. 233 mM sucrose, *W* = 55, *p* = 0.016) **C)** Average isosbestic fluorescence as a function of concentration for the mPFC. Gray dots represent individual animals. We found no significant dependence of fluorescence on concentration (repeated measures ANOVA, *n* = 8, *F* = 1.42 *p* = 0.25). **D)** Same as C for recordings in NAc core. We found no significant dependence of fluorescence on concentration (repeated measures ANOVA, *n* = 9, *F* = 1.93 *p* = 0.12). **E)** Recordings of fluorescence from a single animal around the first lick of 233 mM sucrose showing variability in responses. **F** Perievent average mPFC fluorescence around the beam break crossing that triggers the carousel turn. The light blue region indicates the time during which the carousel turned. Liquid was released at the end of the turn. **G** Same as F for the NAc core.

**Fig. S5.**
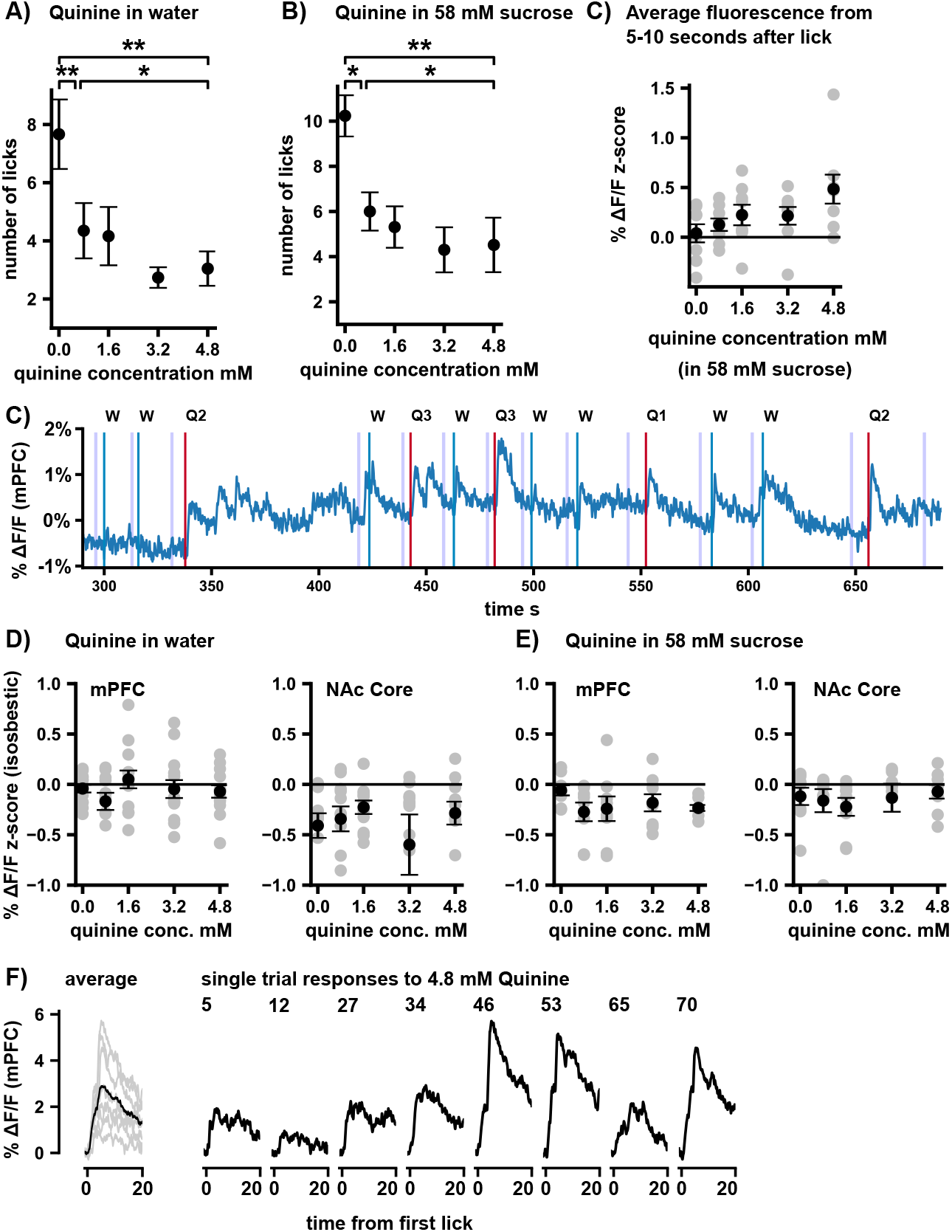
Additional details of recordings during water, quinine, and mixed sucrose and quinine consumption. (* *p <* 0.05, ** *p <* 0.01). **A)** Average number of licks at ports with different concentrations of quinine. Lick rate had a significant dependence on concentration (repeated measures ANOVA, *n* = 24, *F* = 7.63, *p* = 2.4 *×* 10^−5^). We compared lick rates at ports with water and the lowest and highest concentrations of quinine and found that all were significantly different (Wilcoxon sign rank test with Bonferroni correction for multiple comparisons, water vs. 0.8 mM quinine, *W* = 27, *p* = 0.0067; water vs 4.8 mM quinine, *W* = 41, *p* = 0.0033, 0.8 mM quinine vs. 4.8 mM quinine, *W* = 55, *p* = 0.016). **B)** Same as A, but with 58 mM sucrose in all of the solutions. We found a significant dependence of licks at the port with concentration (repeated Measures ANOVA, *n* = 16, *F* = 15.04, *p* = 1.4 *×* 10^−8^). Comparing water and the highest and lowest concentrations of quinine, we found that all were significantly different (Wilcoxon sign rank test with Bonferroni correction for multiple comparisons, water vs. 0.8 mM quinine, *W* = 7, *p* = 0.0017; water vs 4.8 mM quinine, *W* = 14, *p* = 0.01, 0.8 mM quinine vs. 4.8 mM quinine, *W* = 20, *p* = 0.033). **C)** Average fluorescence from 5-10 seconds after the first lick at the port. We found a significant dependence of fluorescence on concentration (repeated measures ANOVA, *F* = 2.9, *p* = 0.040). We also compared pure sucrose with the highest and lowest concentrations of quinine, but did not find any significant pairs (paired *t*-test with Bonferroni correction for multiple comparisons, pure sucrose vs. 0.8 mM quinine, *t* = 1.01, *p* = 1.0; pure sucrose vs 4.8 mM quinine, *t* = 2.38, *p* = 0.15, 0.8 mM quinine vs. 4.8 mM quinine, *t* = 2.30, *p* = 0.16). **D)** Recordings of the z-scored isosbestic fluorescence at different concentrations of quinine. We found no significant dependence of fluorescence on concentration (repeated measures ANOVA; mPFC, *n* = 13, *F* = 1.02, *p* = 0.41; NAc core, *n* = 11, *F* = 0.83, *p* = 0.51). **E)** Same as panel D, but with all solutions containing 58 mM sucrose. We found no significant dependence of fluoresence on concentration (repeated measures ANOVA; mPFC, *n* = 8, *F* = 1.31, *p* = 0.29; NAc core *n* = 8, *F* = 0.80, *p* = 0.53) **F)** Recordings of individual trial responses to licking 4.8 mM quinine from a single animal. Note the large variance in sizes of responses.

**Fig. S6.**
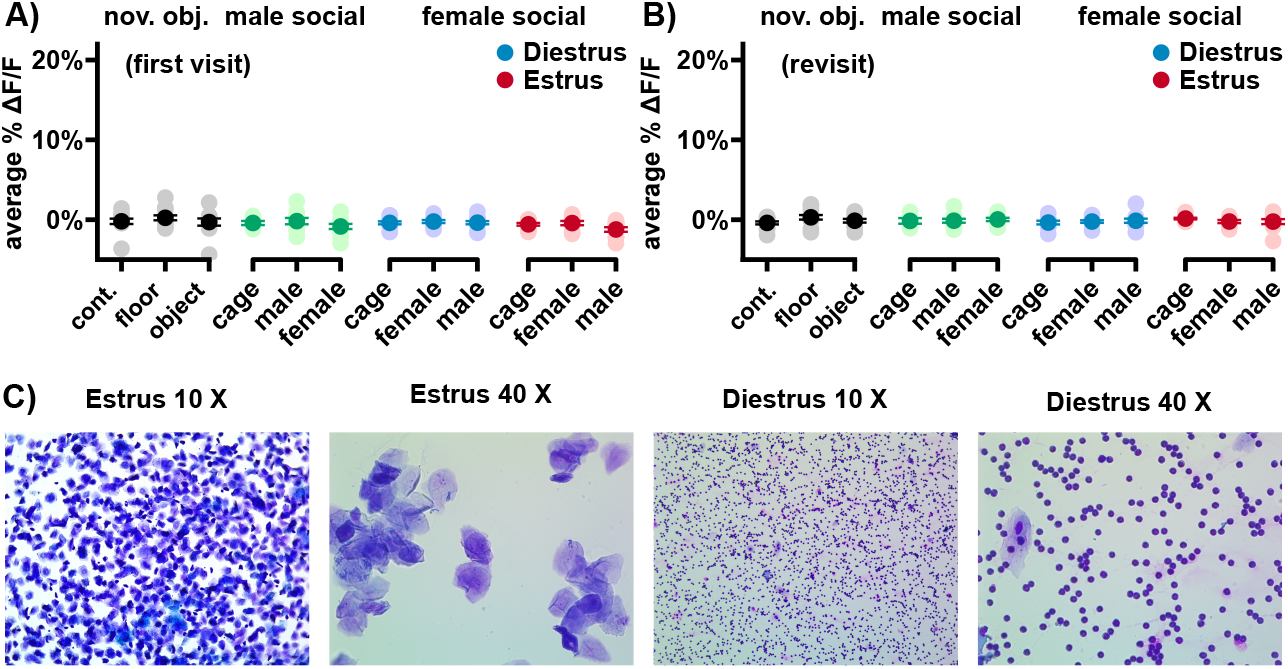
Additional details of the drawer experiment. (* *p <* 0.05, ** *p <* 0.01). **A)** Average mPFC isosbestic fluorescence over the first 5 seconds after a mouse enters the drawer with a stimulus. No significant dependence on stimulus was detected (one-way ANOVA; novel object *n* = 11-12, *F* = 1.14, *p* = 0.56; male social interaction, *n* = 5-11, *F* = 1.58, *p* = 0.45; female social interaction (diestrus), *n* = 9-12, *F* = 0.39, *p* = 0.82; female social interaction (estrus), *n* = 9-11, *F* = 3.78, *p* = 0.15). **B)** Same as panel A for the first visit to the drawer after the stimulus was removed. No significant dependence on stimulus was detected (one-way ANOVA; novel object *n* = 11-12, *F* = 3.38, *p* = 0.18; male social interaction, *n* = 5-10, *F* = 0.28, *p* = 0.86; female social interaction (diestrus), *n* = 9-11, *F* = 0.36, *p* = 0.83; female social interaction (estrus), *n* = 8-11, *F* = 2.85, *p* = 0.24). **C)** Example images of stained vaginal smears used for tracking mouse estrus cycles in estrus (left two panels) and diestrus (right two panels).

## Notes

https://github.com/iellwood/PFCDAAndTotalValence/tree/main

### Competing Interest Statement

The authors have declared no competing interest.

